# Runx1 regulates critical factors that control uterine angiogenesis and trophoblast differentiation during placental development

**DOI:** 10.1101/2023.03.21.532831

**Authors:** Athilakshmi Kannan, Jacob R. Beal, Alison M. Neff, Milan K. Bagchi, Indrani C. Bagchi

## Abstract

During early pregnancy in humans and rodents, uterine stromal cells undergo a remarkable differentiation to form the decidua, a transient maternal tissue that supports the growing fetus. It is important to understand the key decidual pathways that orchestrate the proper development of the placenta, a key structure at the maternal-fetal interface. We discovered that ablation of expression of the transcription factor Runx1 in decidual stromal cells in a conditional *Runx1*-null mouse model (*Runx1*^d/d^) causes fetal lethality during placentation. Further phenotypic analysis revealed that uteri of pregnant *Runx1*^d/d^ mice exhibited severely compromised decidual angiogenesis, and a lack of trophoblast differentiation and migration, resulting in impaired spiral artery remodeling. Gene expression profiling using uteri from *Runx1*^d/d^ and control mice revealed that Runx1 directly controls the decidual expression of the gap junction protein connexin 43 (also known as GJA1), which was previously shown to be essential for decidual angiogenesis. Our study also revealed a critical role of Runx1 in controlling insulin-like growth factor (IGF) signaling at the maternal-fetal interface. While Runx1-deficiency drastically reduced the production of IGF2 by the decidual cells, we observed concurrent elevated expression of the IGF-binding protein 4 (IGFBP4), which regulates the bioavailability of IGFs thereby controlling trophoblast differentiation. We posit that dysregulated expression of GJA1, IGF2, and IGFBP4 in *Runx1*^d/d^ decidua contributes to the observed defects in uterine angiogenesis, trophoblast differentiation, and vascular remodeling. This study therefore provides unique insights into key maternal pathways that control the early phases of maternal-fetal interactions within a critical window during placental development.

**Significance:** A clear understanding of the maternal pathways that ensure coordination of uterine differentiation and angiogenesis with embryonic growth during the critical early stages of placenta formation still eludes us. The present study reveals that the transcription factor Runx1 controls a set of molecular, cellular, and integrative mechanisms that mediate maternal adaptive responses controlling uterine angiogenesis, trophoblast differentiation, and resultant uterine vascular remodeling, which are essential steps during placenta development.

## INTRODUCTION

During early pregnancy, the endometrium transitions from a functionally non-receptive to a receptive state that allows embryo implantation. This critical transformation, orchestrated by the ovarian steroid hormones, must be synchronized with embryonic development to ensure maximal reproductive success (1–4). To enable this synchronization, an intricate maternal-fetal dialogue has evolved that allows the developing embryo and the endometrium to be in constant communication with each other, acting in concert to direct the events that lead to the formation of a functional placenta (5, 6).

In humans and rodents, as embryo implantation is initiated, the uterus undergoes a dramatic transformation to form the decidua, a transient stroma-derived secretory tissue that surrounds the growing fetus (1–3). Decidual cells produce and secrete various paracrine factors that act on neighboring cells within the uterine milieu to control diverse functions, including the formation of an extensive vascular network that supports embryo development (7, 8). Proper differentiation and migration of the trophoblast cells, critical for forming functional placenta, are also influenced by as yet unknown factors secreted by the decidual cells (5, 6). Any disruption in these processes is likely to hinder proper placenta development, resulting in early pregnancy failure and vascular impairments seen in gestational diseases such as recurrent miscarriage, preeclampsia, and intrauterine growth restriction (9–12).

The mechanisms by which maternal factors promote embryonic growth and survival during the time between embryo implantation and placental development remain an important unresolved question in biology. Although several factors controlling uterine proliferation and differentiation have been identified by murine transcriptomics and by using mutant mouse models that show impairments in early phases of implantation, a clear understanding of the molecular pathways that ensure coordination of the endometrial differentiation and angiogenesis with embryonic growth during more advanced phases of implantation and decidualization, particularly during the critical early stages of placentation, still eludes us. This is primarily due to a lack of experimental models addressing the mechanisms by which maternal factors control placenta development.

In this study, we have developed and analyzed a mouse model *Runx1*^d/d^ in which the gene encoding the RUNT family transcription factor, Runx1, is conditionally deleted in endometrial stromal cells. We focused on this factor because it is expressed in decidual stromal cells following implantation and is a downstream target of regulation by bone morphogenetic protein 2 (BMP2) which is a critical regulator of implantation (13, 14). The *Runx1*^d/d^ female mice displayed no apparent defects in implantation or decidualization but exhibited severely reduced fertility, arising from placental abnormalities. Strikingly, most embryos in *Runx1* ^-^null uteri failed to grow as the placenta begins to develop. Phenotypic analysis of uteri lacking *Runx1* revealed highly compromised decidual angiogenesis and impairments in trophoblast differentiation and migration into the decidua. The *Runx1*^d/d^ mouse model therefore presents a unique opportunity to test the hypothesis that maternal decidual deficiencies contribute to placental defects and pregnancy failure. Here, we will describe our results uncovering the cell-cell signaling mechanisms that mediate maternal adaptive responses controlling uterine angiogenesis and trophoblast differentiation and invasion, which are essential steps during placenta development.

## RESULTS

### *Runx1* is induced downstream of BMP2 signaling in mouse endometrial stromal cells during decidualization

In mice, implantation is initiated four days after fertilization, when the blastocysts attach to the uterus (15). Previous studies in the DeMayo laboratory and our laboratory revealed that bone morphogenetic protein 2 (BMP2) is an essential regulator of implantation in mice and it controls differentiation of endometrial stromal cells during early pregnancy (13, 14, 16). Gene expression profiling experiments revealed *Runx1* as a potential downstream target of BMP2 in pregnant uterus (13). To confirm this finding, we employed a primary culture system in which undifferentiated stromal cells isolated from pregnant mouse uteri (preimplantation, day 4) undergo decidualization *in vitro* in the presence or absence of BMP2. As shown in Fig. 1A, a marked increase in the expression of *Runx1* transcripts was observed in mouse endometrial stromal cells within 24 h after the addition of BMP2, establishing that Runx1 is indeed a downstream target of regulation by BMP2 during decidualization.

**Figure. 1.**
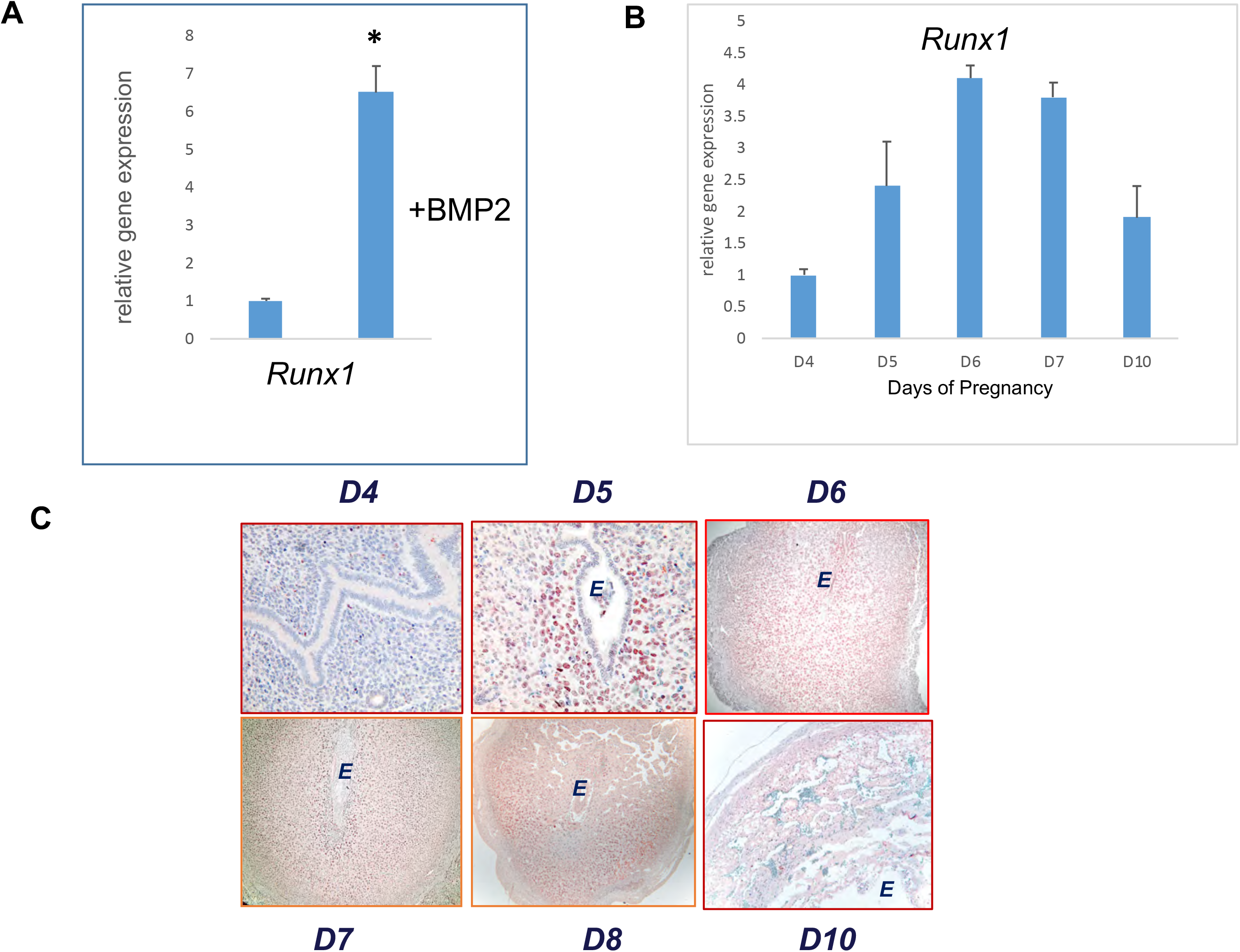
Runx1 is induced in the uterus during early pregnancy. **A.** The primary cultures of mouse endometrial stromal cells were treated with or without recombinant BMP2 for 24 h. The cells were then lysed and total RNA was isolated and real-time PCR was performed to analyze the levels of *Runx1*. The relative levels of gene expression were determined by setting the expression level of sample without BMP2 treatment to 1.0 (n=3). *Rplp0,* encoding a ribosomal protein, was used to normalize the level of RNA. *P < 0.05. **B**. qPCR was performed to monitor the expression of *Runx1* mRNA in uteri on days 4 to 10 of gestation. The relative levels of gene expression on different days of pregnancy were determined by setting the expression level of *Runx1* mRNA on day 4 of pregnancy at 1.0. *Rplp0,* encoding a ribosomal protein, was used to normalize the level of RNA. Data represent mean ± SEM from three separate samples. **C.** Uterine sections on days 4 (panel a), 5 (panel b), 6 (panel c), 7 (panel d), 8 (panel e), and 10 (panel f) of pregnancy were subjected to immunohistochemical analysis using anti-RUNX1 antibody. Data are representative images from n=4. E indicates an embryo.

We next examined the spatio-temporal profiles of mRNA and protein corresponding to *Runx1* in the mouse uterus on days 4 to 10 of pregnancy using real-time PCR and immunohistochemistry (IHC). As shown in Fig. 1B, *Runx1* mRNA expression was induced on day 5 of pregnancy, increased further on days 6 and 7, overlapping the decidualization period, and continued on day 10, although at a reduced level, following initiation of placentation. Consistent with the mRNA profile, Runx1 protein expression (Fig. 1C) was undetectable in uterine sections obtained from day 4 pregnant mice in the period preceding implantation (panel a). Distinct Runx1 immunostaining was visible in the stromal cells surrounding the implanted embryo in the uterine sections of mice on day 5 of pregnancy (panel b). Widespread Runx1 expression was observed in the decidual stromal cells surrounding the implanted embryo as pregnancy progressed to days 6 to 8 (panels c, d, and e). The expression of Runx1 protein then declined by day 10 of gestation (panel f), suggesting a possible functional role of this factor during decidualization and early placentation in pregnant mouse.

### Conditional deletion of *Runx1* in the uterus leads to severe infertility

Previous studies showed that the *Runx1*-null embryos die at ∼day 12.5 due to defects in hematopoiesis (17, 18). To gain insight into the role of *Runx1* in adult mice during pregnancy, we therefore crossed mice harboring the “floxed” *Runx1* (*Runx1^f/f^*) locus with the progesterone receptor (Pgr)-Cre mice to create *Runx1^d/d^* mice (19). The *Runx1*^d/d^ mice thus generated are heterozygous for the Pgr-Cre and homozygous for the “floxed” *Runx1* conditional allele. We assessed the extent of deletion of *Runx1* in the uteri of *Runx1^d/d^* mice by qPCR and immunofluorescence (Fig. 2). Our results showed greatly reduced expression of *Runx1* transcripts, but not *Pgr* transcripts, in the uteri of *Runx1^d/d^* mice on day 7 of pregnancy, indicating efficient ablation of *Runx1* gene in the uteri of these mice (Fig. 2A). Consistent with the loss of *Runx1* mRNA expression, we observed a marked decline in the level of Runx1 protein in *Runx1^d/d^* uteri on day 7 of gestation (Fig. 2B), indicating efficient ablation of Runx1 expression in the uterine stromal cells.

**Figure 2.**
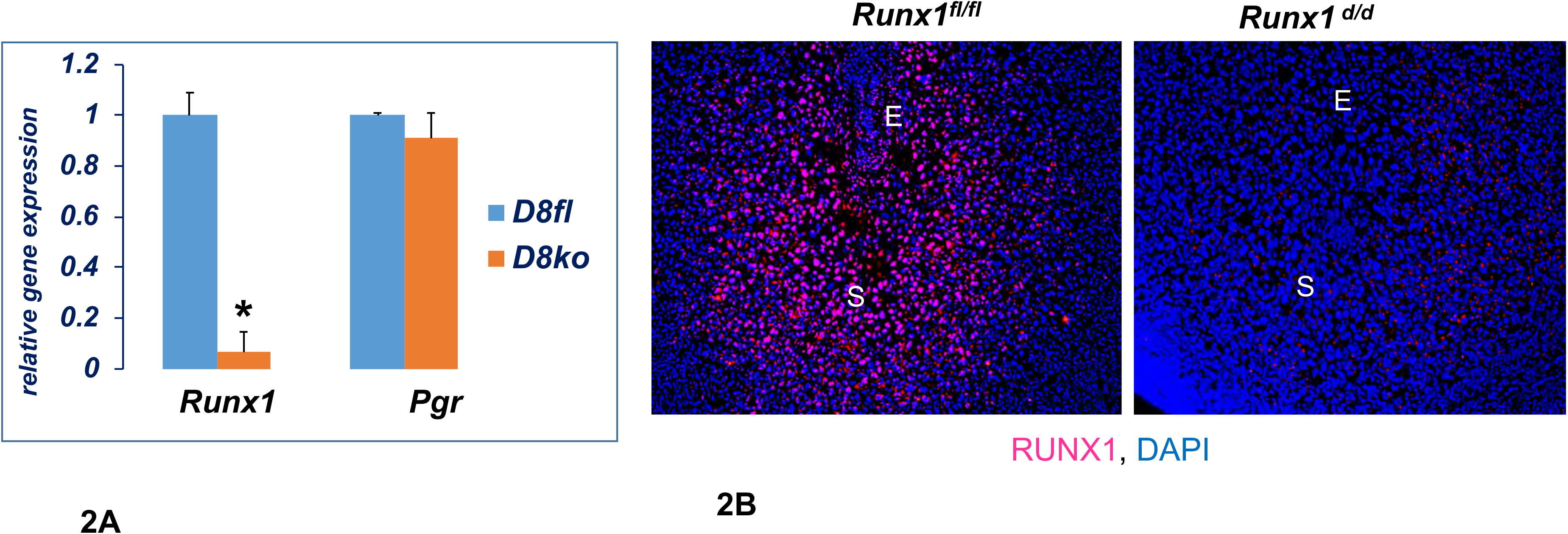
Loss of Runx1 expression in the uterus of *Runx1^d/d^* mice. **A.** Uterine RNA was purified from *Runx1^f/f^* and *Runx1^d/d^* mice on day 8 of pregnancy and analyzed by qPCR. Relative levels of *Runx1* mRNA and *Pgr* mRNA expression in uteri of *Runx1^d/d^* mice are compared to those in *Runx1^f/f^* control mice. Data represent mean ± SEM from three separate samples. Asterisk indicates statistically significant differences (\**P < 0*.05). **B.** Uterine sections obtained from day 8 pregnant *Runx1^f/f^* (left panel) and *Runx1^d/d^* (right panel) mice were subjected to IF using anti-Runx1 antibody. Note the lack of Runx1 immunostaining in uteri of the mutant mice. E and S indicate embryo and stroma, respectively.

We then performed a six-month breeding study in which the *Runx1*^d/d^ female mice were crossed with wild-type males. These females exhibited severe fertility defects, showing more than 80% reduction in the total number of pups born per *Runx1*^d/d^ female compared with a *Runx1*^f/f^ female (Fig. S1). The few pups that were born to the *Runx1*^d/d^ females had severely low birth weights and died within 48h of birth. It is important to point out that the *Runx1* allele is intact in the embryos resulting from crosses of *Runx1*^d/d^ females with wild-type males. Thus, the phenotypic defects observed during pregnancy in *Runx1*^d/d^ mice are not due to an intrinsic lack of this gene in the embryos but arises from *Runx1* deficiency in the maternal tissue.

We next investigated whether the infertility of *Runx1^d/d^* females was due to an ovarian defect. An analysis of the ovulation in *Runx1^f/f^* and *Runx1^d/d^* females revealed no significant difference in the number of released oocytes (Fig. S2A) between the two genotypes. Histological examination of the ovaries confirmed the presence of corpora lutea in both types of mice (Fig. S2B). The serum levels of progesterone and 17*β*-estradiol were unaltered in pregnant *Runx1*^d/d^ mice on day 8 of gestation (Fig. S2C), so there is no evidence that steroidogenesis in the corpus luteum is compromised in *Runx1*^d/d^ mice during pregnancy. Collectively, these results indicated that the infertility of *Runx1^d/d^* females is not due to impairment in the hypothalamic-pituitary-ovarian axis but is likely due to defective uterine function.

### Embryo attachment and early stages of decidualization are unaffected in *Runx1^d/d^* mice

Gross examination of uterine morphology revealed apparently normal embryonic implantation sites in *Runx1^f/f^* and *Runx1^d/d^* uteri on day 6 of pregnancy (Fig. 3A), implying embryo attachment was unaffected by the absence of Runx1. We then examined the expression of a panel of factors, *Pgr*, *Bmp2,* and *C/ebp1*, which are known regulators of decidualization in mice (13, 14, 20, 21). Our studies showed that the expression of *Pgr*, *Bmp2*, and *C/ebpb* mRNAs remained unaffected by the loss of uterine Runx1 (Fig. 3B). Consistent with these observations, the expression of alkaline phosphatase, a well-known biomarker of decidualization was comparable in the uterine sections of *Runx1^f/f^* and *Runx1^d/d^* mice on day 7 of pregnancy, indicating that at least certain aspects of the decidualization process progress normally in *Runx1^d/d^* uteri (Fig. 3C).

**Figure 3.**
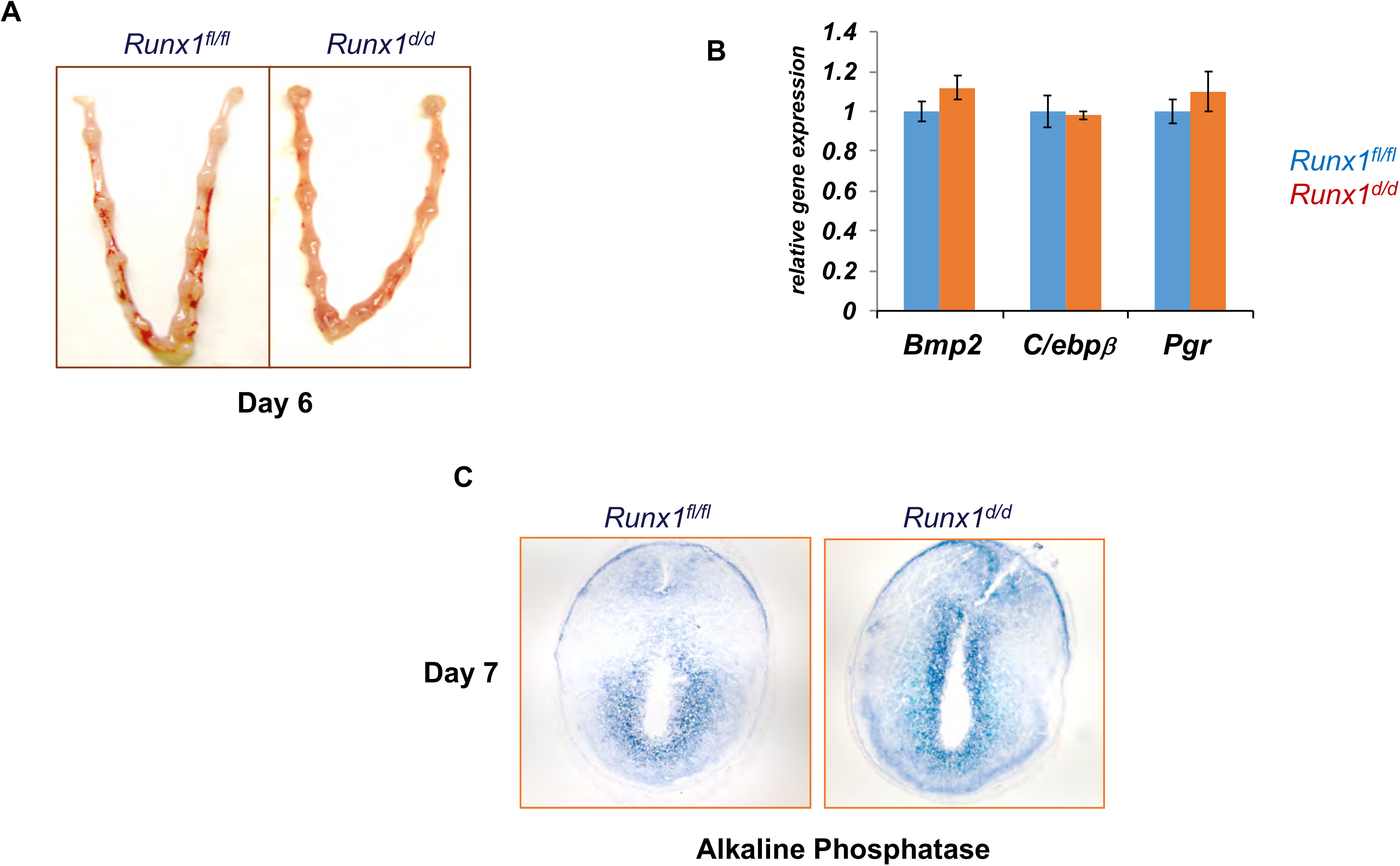
Early implantation is unaffected in *Runx1^d/d^* mice. **A.** Gross morphology of *Runx*1*^f/f^* and *Runx*1*^d/d^* uteri at day 6 of gestation. **B.** Comparable expressions of various markers of decidualization in *Runx*1*^f/f^* and *Runx*1*^d/d^* uteri. Total RNA was isolated from uteri on day 6 of pregnancy and qPCR analysis was performed using primers specific for *Bmp2* and *Cebpb* and *Pgr*. **C.** Uterine sections from *Runx*1*^f/f^* and *Runx*1*^d/d^* mice on day 7 of pregnancy were subjected to alkaline phosphatase activity (n=3).

### Decidual angiogenesis is impaired in *Runx*1*^d/d^* mice

Although no apparent functional abnormality was detected in pregnant *Runx1^d/d^* uteri up to day 8 of gestation, we observed distinct signs of hemorrhage and embryo resorption in these uteri starting on day 10 and by day 15 most of the implantation sites had regressed (Fig. 4). When the decidual swellings were dissected, they contained underdeveloped placenta with resorbed embryos (Fig. 4, bottom, right). We performed PCR-based genetic sex determination assays, which indicated that sexual dimorphism does not exist in placental/fetal development in *Runx1^d/d^* mice (22).

**Figure 4.**
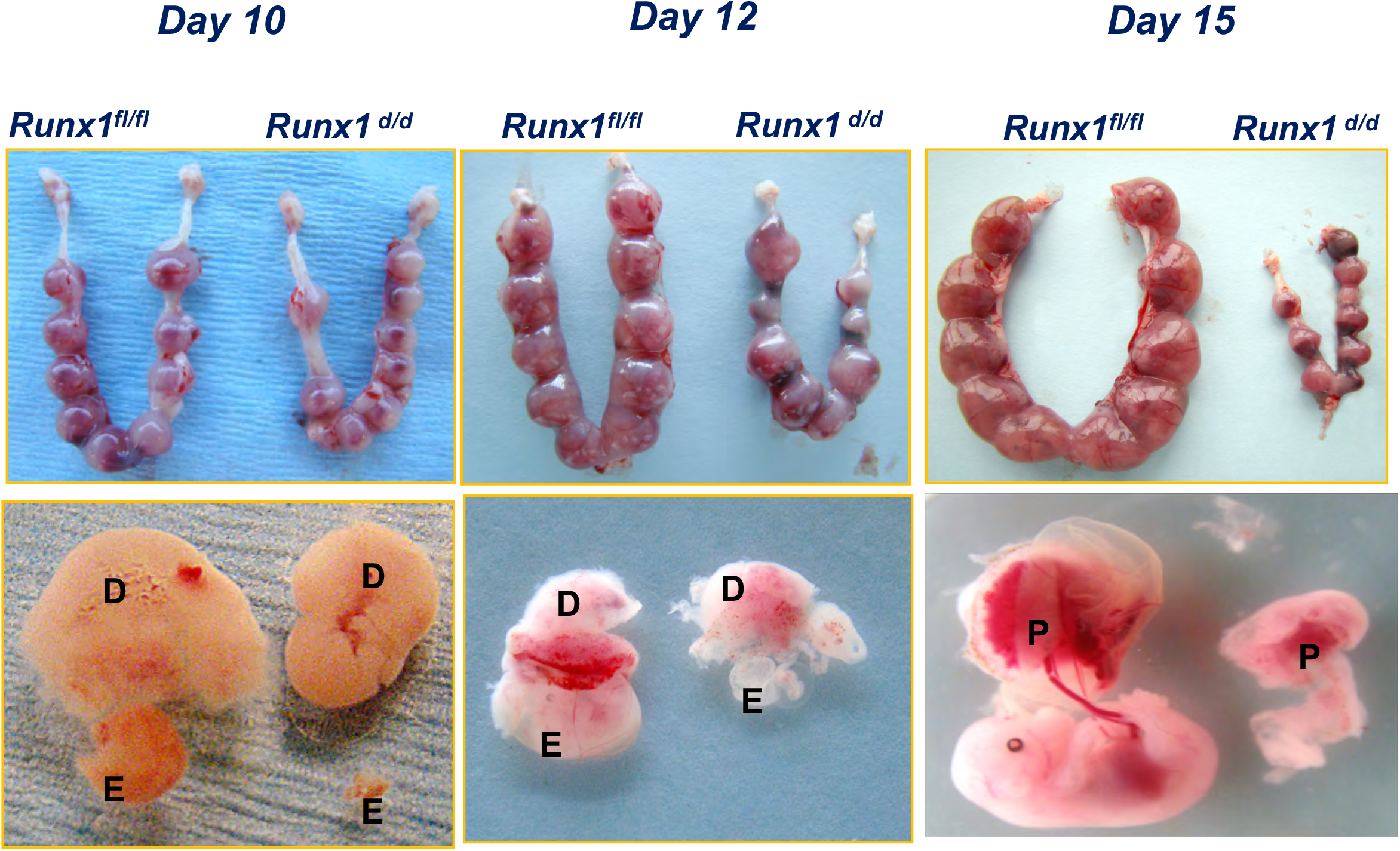
Pregnancy failure in *Runx*1*^d/d^* mice in mid gestation. **Upper.** Gross morphology of *Runx*1*^f/f^* and *Runx*1*^d/d^* uteri at days 10, 12 and 15 of gestation (n=5/genotype on each day). **Lower.** Dissection of implantation sites revealed underdeveloped placenta and degenerating embryos in *Runx1^d/d^*. E, D, and P indicate embryo, decidua and placenta, respectively.

We next examined the development of vascular networks in pregnant uteri of *Runx1^d/d^* mice by employing immunofluorescence using an antibody against platelet/endothelial cell adhesion molecule 1 (PECAM1), a marker of endothelial cells. Uterine sections of the control *Runx1^f/f^* mice on day 8 of pregnancy exhibited a well-developed vascular network that spreads throughout the decidual bed surrounding the implanted embryo (Fig. 5A, panel a). In contrast, the PECAM1 immunostaining was markedly reduced in pregnant *Runx1^d/d^* mice, indicating a severe impairment in decidual endothelial network formation (Fig. 5A, panel b). This reduced angiogenesis was associated with considerable hemorrhagic activity in the implantation chambers of *Runx1^d/d^* mice. During early pregnancy, Runx1 is expressed in endometrial stromal cells but not in luminal or glandular epithelium. Also note that *Runx1* is not expressed in uterine endothelial cells, as dual IF indicates non-overlapping expression of Runx1 (green) and PECAM (red) (Fig. 5A, panels c and d). Therefore, the observed defect in angiogenesis in *Runx1*^d/d^ uteri is not due to *Runx1* deficiency in epithelial or endothelial cells but a result of excision of this gene in the uterine stromal cells. Collectively, these results indicated that the reproductive defect in the *Runx1*^d/d^ female is of uterine stromal origin and manifests within a critical time window between implantation and initial phases of placentation.

**Figure 5.**
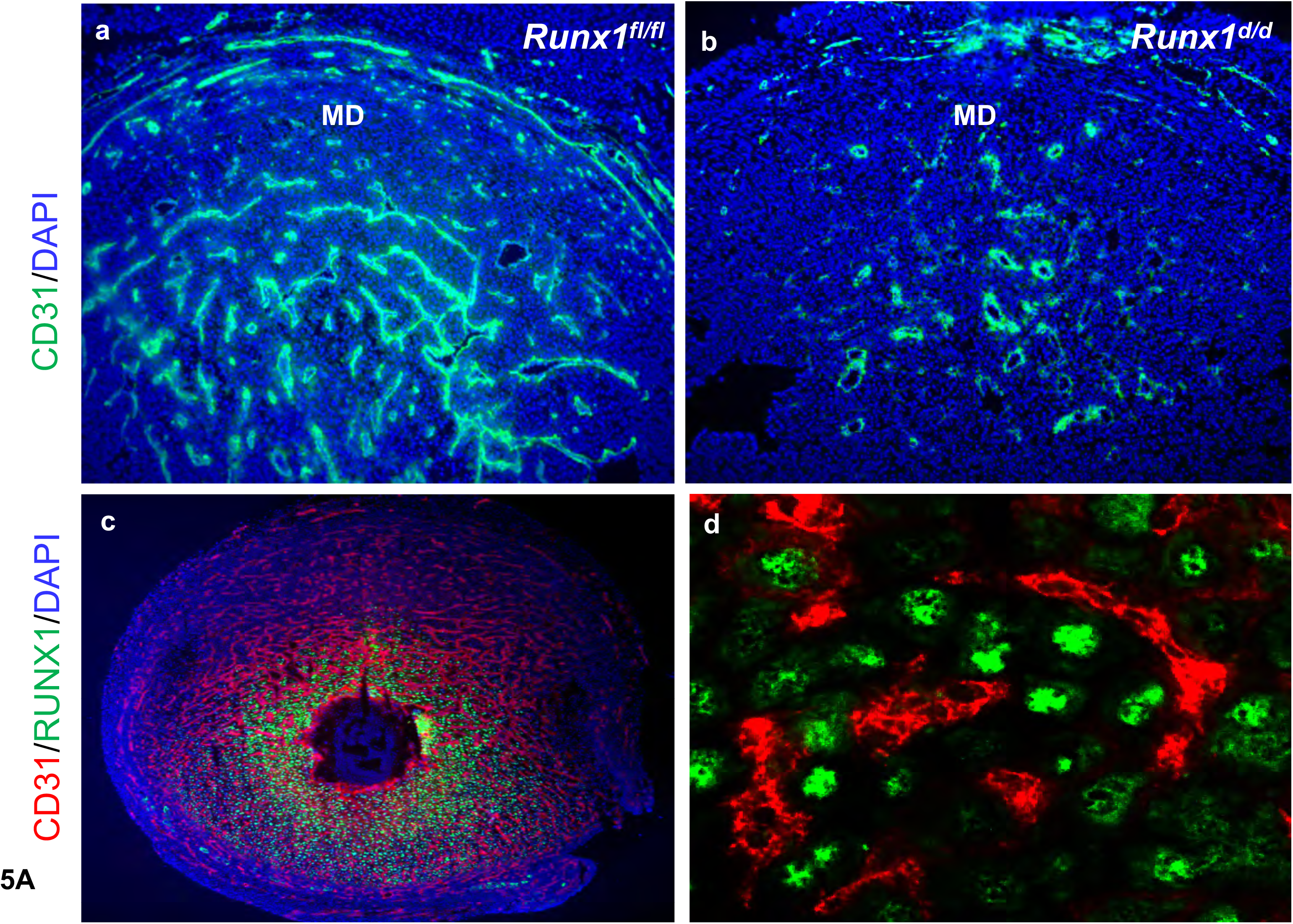
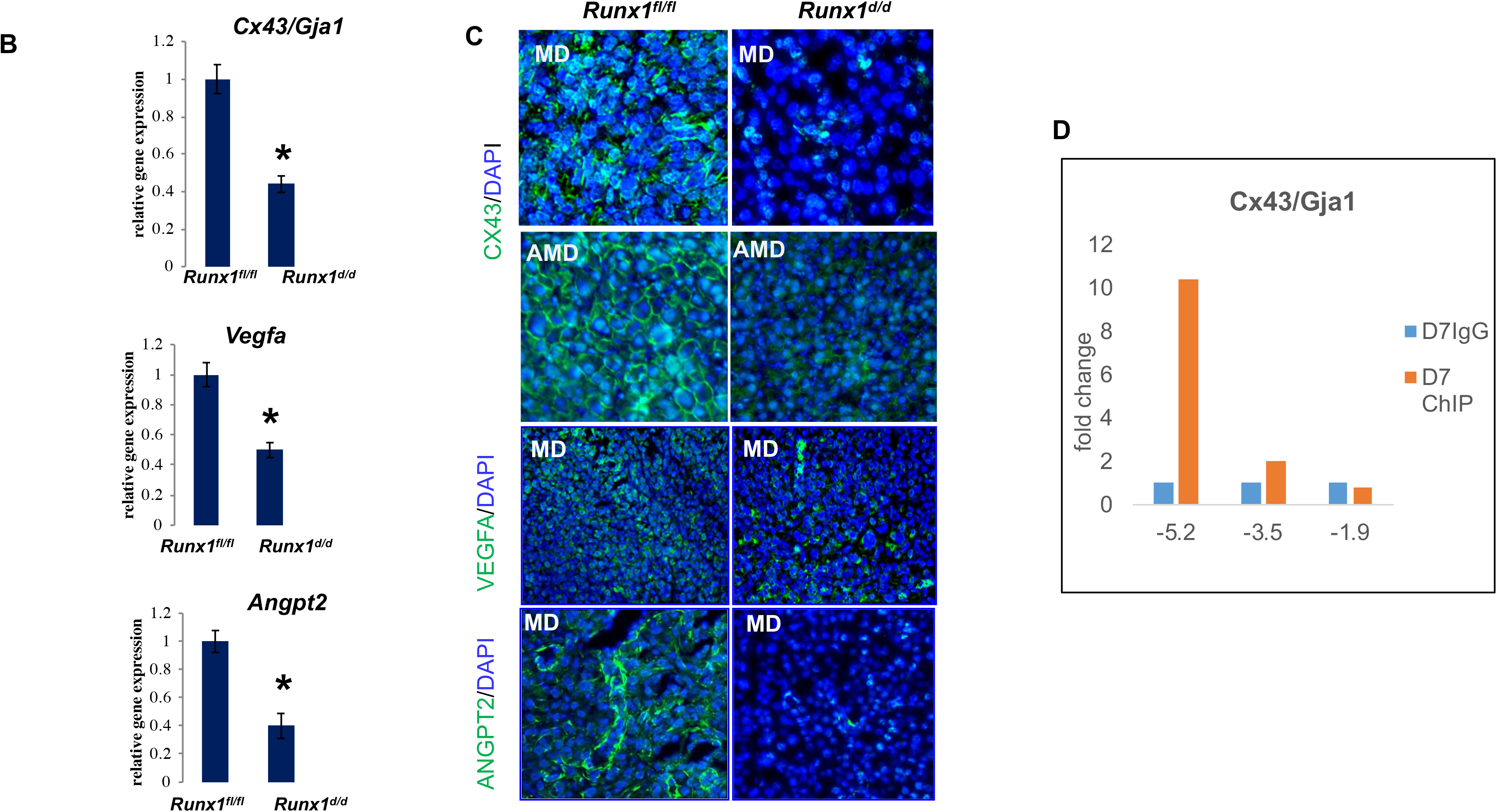
Pregnancy failure in *Runx*1*^d/d^* mice is associated with lack of angiogenesis. **A. Upper:** Uterine sections of *Runx*1*^f/f^* (panel a) and *Runx*1*^d/d^* (panel b) mice on day 8 (n=4) were subjected to IF staining with CD31 antibody. Representative images are shown. MD denote mesometrial decidua. **Lower:** Dual IF analysis in *Runx1*^f/f^ uteri shows that RUNX1 protein (green) is expressed in the stromal cells, but not in the endothelial cells (PECAM, red) (panel c). Panel d represents magnified images of stromal and endothelial cells. **B.** qPCR was performed to analyze the expression of angiogenic factors, *Cx43*/*Gja1*, *Vegfa* and *Angpt2* in uteri of *Runx*1*^f/f^* and *Runx*1*^d/d^* mice on day 8 of pregnancy. Data represent mean ± SEM from four separate samples and were analyzed by *t*-test. Asterisks indicate statistically significant differences (\**P < 0*.05). **C.** Uterine sections of *Runx*1*^f/f^* and *Runx*1*^d/d^* mice on day 8 were subjected to IF staining with CX43/GJA1, VEGFA, and ANGPT2 antibodies. Data are representative images from n=3. AMD and MD denote antimesometrial decidua and mesometrial decidua, respectively. **D.** Stromal cells from day 7 pregnant mouse uteri was analyzed by chromatin immunoprecipitation (ChIP) using Runx1 antibody (orange) and control antibody (blue). Chromatin enrichment was quantified by real-time PCR using primers flanking the RUNT binding element in the *Cx43*/*Gja1* promoter. Enrichments were normalized to 1% of input DNA. Runx1 occupied the -5.2 kb region of the *Cx43*/*Gja1* gene. The experiment was repeated twice; representative data are shown.

### Runx1 controls the expression of Cx43 which is essential for uterine angiogenesis

To investigate the biological pathways affected by *Runx1* deletion in the uterus, we performed gene expression profiling. Microarray analysis of decidual cells isolated from *Runx1^fl/fl^* and *Runx1^d/d^* uteri on day 10 of pregnancy revealed downregulation of mRNAs corresponding to many genes among which those controlling vascular development/remodeling, metabolic processes, and cell proliferation/differentiation. Because of the vascular defect and hemorrhage in *Runx1^d/d^* uteri, we focused on the angiogenesis-related pathways that are downregulated in the absence of Runx1. We confirmed that the expression of several factors, including *Cx43/Gja, Vegfa*, and *Angpt2,* were markedly downregulated in *Runx1^d/d^* uteri compared to *Runx1^f/f^* uteri (Fig. 5B). Consistent with the mRNA profiles, we observed a marked downregulation of these proteins in *Runx1^d/d^* uteri (Fig. 5C). We took a particular interest in Cx43, since we had shown previously that conditional deletion of the *Cx43/Gja* gene in *Cx43^d/d^* mice did not affect the hypothalamo-pituitary-ovarian axis, implantation, or the myometrial function during early or mid-pregnancy, but led to a striking impairment in decidual angiogenesis (23). Our studies had previously shown that Cx43 is expressed in the endometrial stromal cells but not in the endothelial cells in the decidua (23). Similarly, in Fig. 5A, we show that Runx1, which regulates Cx43 expression, is expressed exclusively in the stromal cells but not in the endothelial cells. The localization of both Runx1 and Cx43 in stromal but not in endothelial cells indicates that the angiogenic defects observed in *Runx1^d/d^* and *Cx43^d/d^* uteri during pregnancy are not due to deletion of these genes in the endothelial cells but rather in the stromal cells (23). The stromal loss of Cx43, which disrupts the gap junctions between adjacent decidual cells, resulted in impaired angiogenesis, leading to the arrest of embryonic growth by gestation day 10.

We next investigated whether *Cx43/Gja* is a direct target of Runx1 in endometrial stromal cells. Bioinformatic analyses revealed the presence of putative Runx1 binding sites in the 5’-flanking region of the mouse *Cx43/Gja* gene. We then performed chromatin immunoprecipitation (ChIP), using stromal cells isolated from mouse uteri on day 7 of pregnancy when ample Cx43 expression is observed in the decidua. Our results showed that Runx1 binds robustly to a consensus binding site at the -5.2 kb upstream region of the mouse *Cx43/Gja* gene and rather weakly at other putative sites at the -3.5 kb and -1.9 kb regions (Fig. 5D). Our results suggest that *Cx43/Gja* gene is a direct target of regulation by Runx1. Consistent with this finding, as pregnancy progresses from days 7 to 10, the expression of both Runx1 and Cx43 is maintained in the decidua and Runx1-Cx43 pathway presumably directs the expression of downstream angiogenic networks in this tissue (23).

### Abnormal trophoblast differentiation and migration in *Runx1^d/d^* uteri

Since *Runx1*-null uteri exhibited embryo resorption at the onset of placentation, we performed in-depth analyses of uterine sections of *Runx1^fl/fl^* and *Runx1^d/d^* mice on day 10 of gestation. Immunofluorescence analyses using antibodies against PL1, a trophoblast giant cell (TGC) specific marker (24–26), revealed that the placentas of *Runx1^f/f^* mice, as expected, displayed normal characteristics of one to two layers of TGCs at the maternal-fetal interface on day 10 (Fig. 6A). In contrast, we observed multiple layers of TGCs at the maternal-fetal interface in *Runx1^d/d^* animals (Fig. 6A), indicating dysregulated expansion of the TGCs.

**Figure 6.**
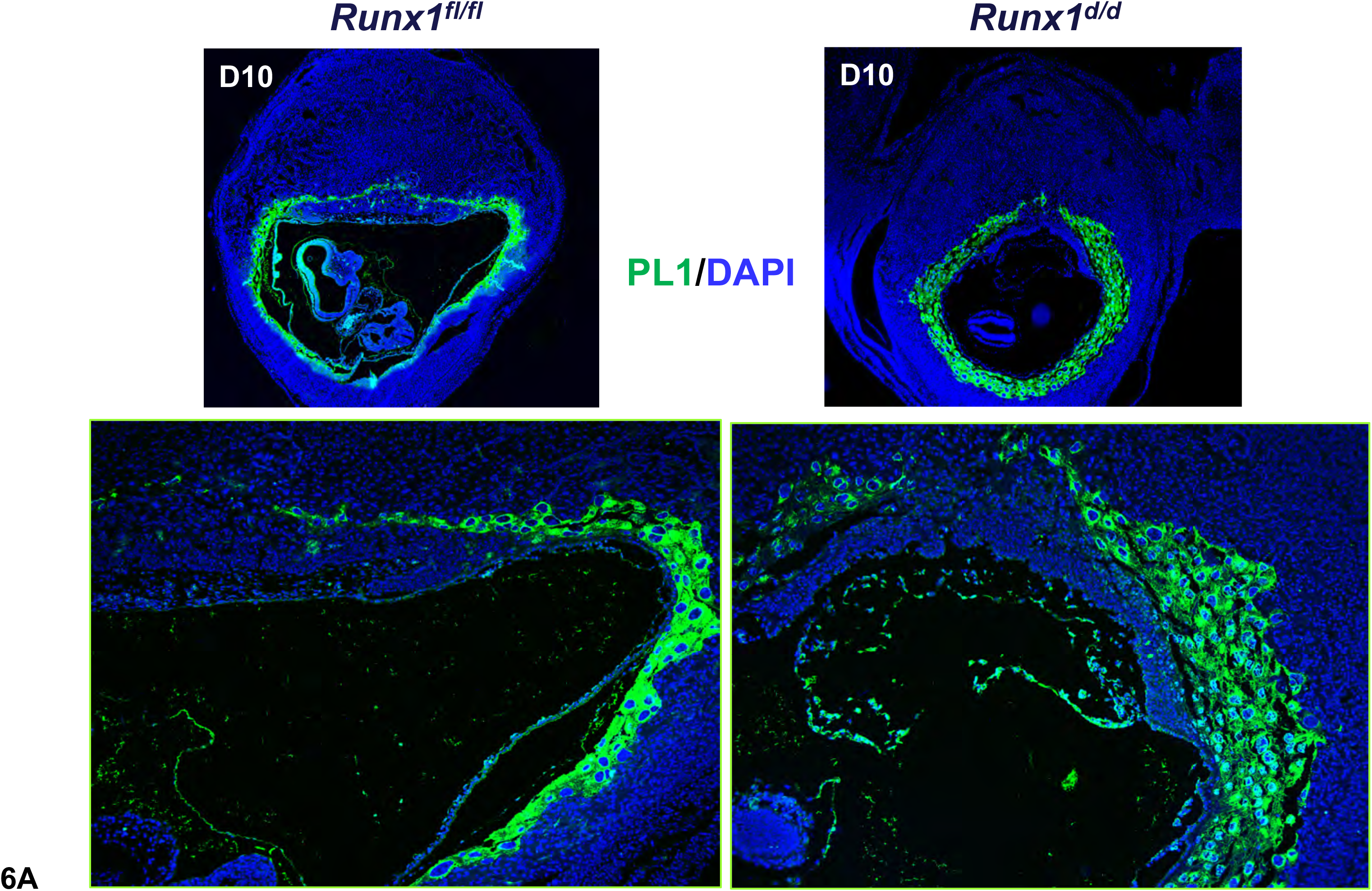
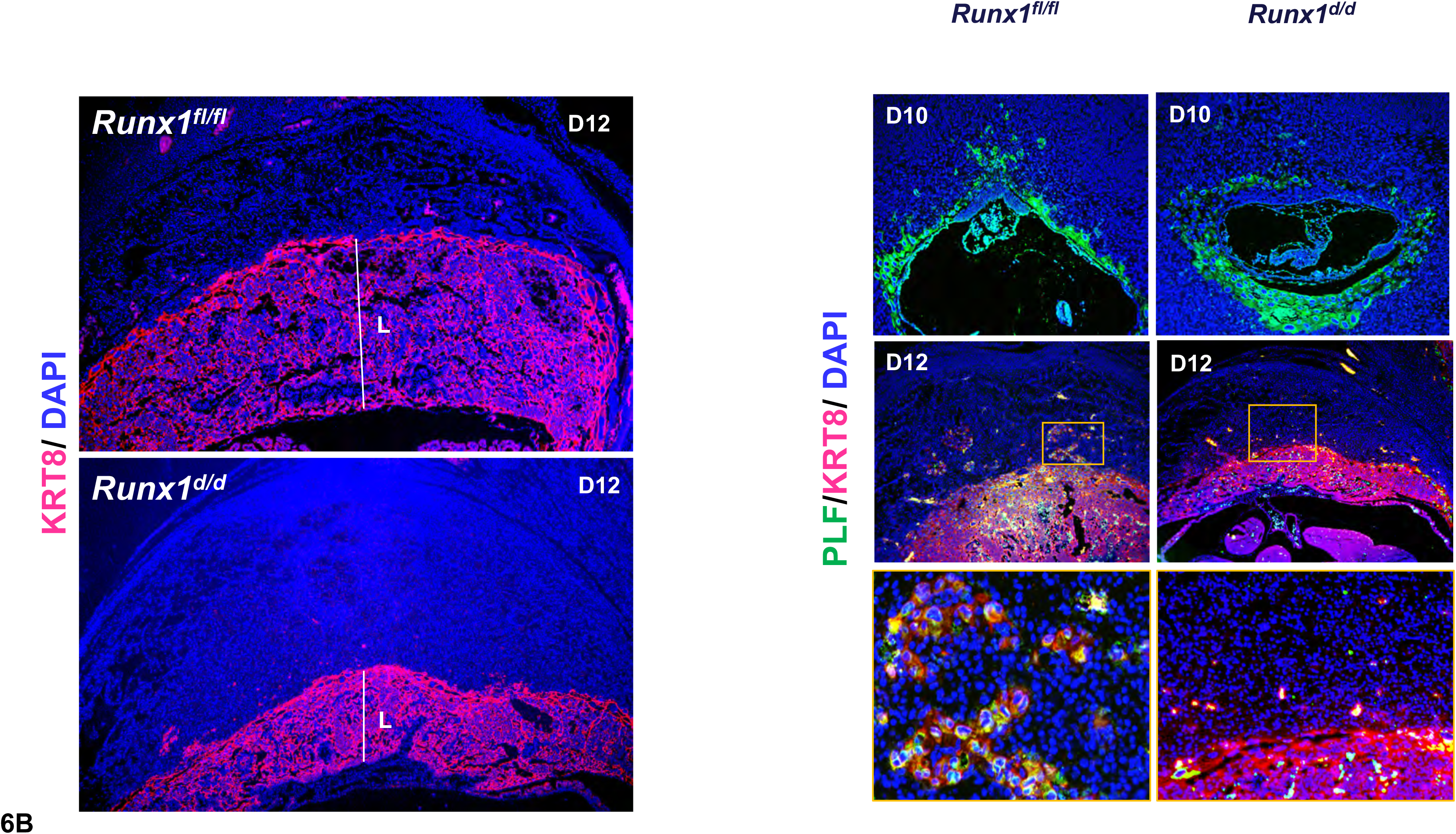
Abnormal trophoblast proliferation, differentiation and disorganized placentation in *Runx*1*^d/d^* mice. **A.** Increased population of trophoblast giant cells (TGCs) in the placentae of *Runx*1*^d/d^* mice. Uterine sections from *Runx1^f/f^* and *Runx1^d/d^* mice on day 10 of pregnancy were subjected to IF using PL1. Lower panels indicate magnified images. **B. Left.** Uterine sections from *Runx1^f/f^* and *Runx1^d/d^* mice on day 12 of pregnancy were subjected to IF using KRT8 antibody. **B. Right.** Uterine sections from *Runx1^f/f^* and *Runx1^d/d^* mice on day 10 and 12 of pregnancy were subjected to IF staining using PLF and KRT8 antibodies. Boxed regions indicating magnified images are shown in the lower panels. Data are representative images from n=3.

With the progression of gestation, trophoblast cells undergo differentiation and, along with fetal blood vessels, form extensive villous branching to create a densely packed structure called the labyrinth (27). Further, a subpopulation of TGCs differentiate into invasive TGCs or spongiotrophoblasts, which are indicated by the expression of their biomarker PLF (24–27). These invasive spongiotrophoblasts migrate into the decidua to reach spiral arteries and dilate them for much needed nutritional support for the growing embryo. Interestingly, our studies revealed that as pregnancy progressed to day 12, the *Runx1^d/d^* placentas appeared to be disorganized, lacking properly formed layers, including a significantly smaller labyrinth (Fig. 6B, left). We also noted with interest a distinct difference in the spatial migration of the spongiotrophoblast cells in the placentas of *Runx1^d/d^* mice compared to *Runx1^fl/fl^* controls (Fig. 6B, right). As expected, PLF positive invasive trophoblast cells were observed in the *Runx1^fl/fl^* decidua on day 10 or 12 of pregnancy (Fig. 6B). In contrast, *Runx1^d/d^* females exhibited curtailed migration of spongiotrophoblasts in the decidua, indicating impaired differentiation of trophoblast cells into an invasive phenotype, leading to restricted trophoblast invasion into *Runx1^d/d^* decidua.

### Lack of Spiral Artery Modification in *Runx1^d/d^* mice

Ample evidence exists that proper trophoblast differentiation and migration into the decidua are functionally linked to efficient spiral artery modifications during placentation. We therefore investigated whether spiral artery modification is indeed impacted by impaired trophoblast differentiation in *Runx1^d/d^* females. Dual immunofluorescence staining using antibodies specific for smooth muscle actin (α-SMA), a marker for arterial smooth muscle cells, and for cytokeratin 8 as a trophoblast marker, revealed arterial dilation in *Runx1^fl/fl^* decidua as indicated by the presence of trophoblast cells and the absence of smooth muscle cells in the spiral arteries (Fig.7A). However, the trophoblasts did not reach the spiral arteries to modify these vessels in *Runx1^d/d^* decidua.

**Figure 7.**
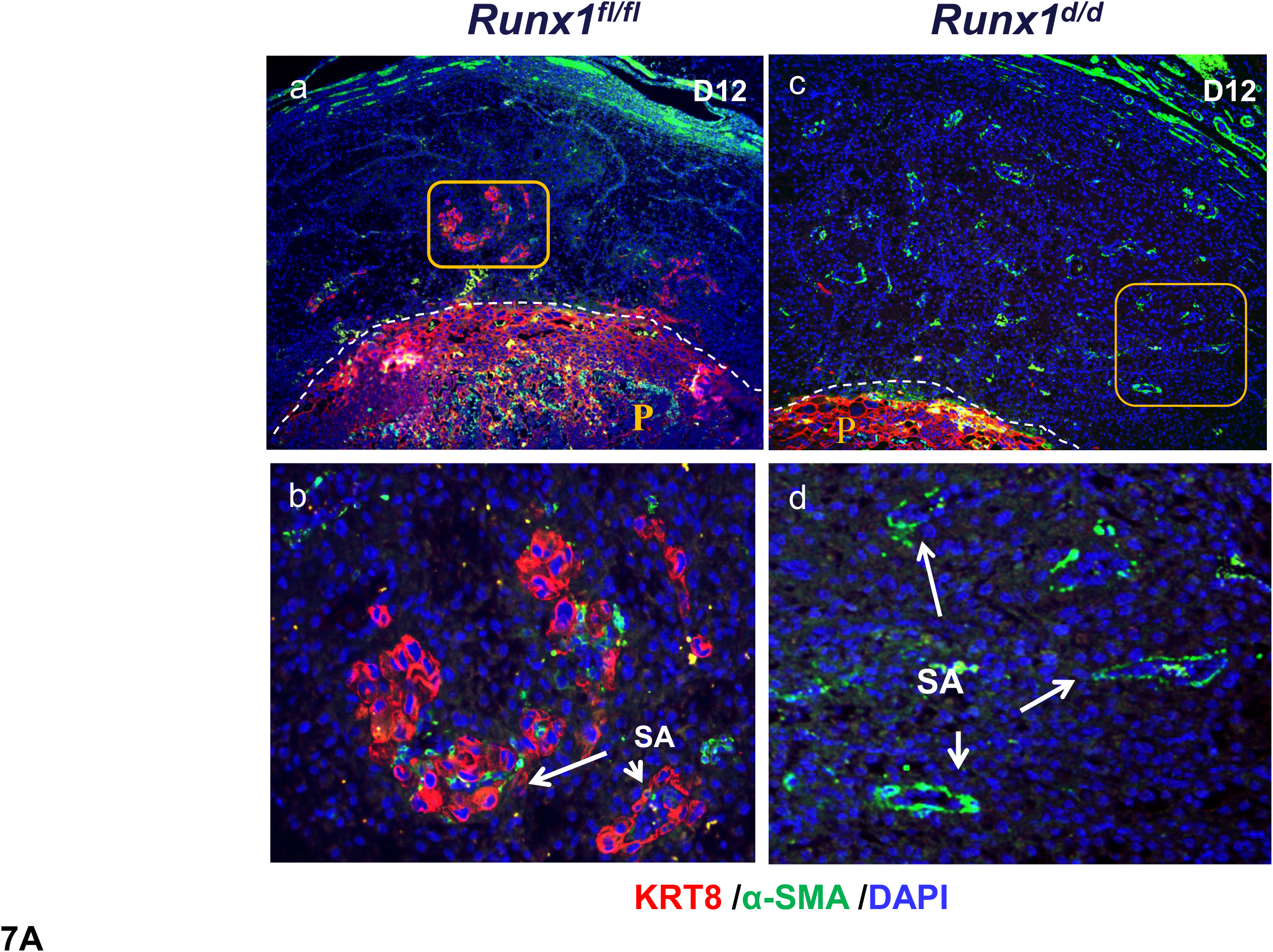
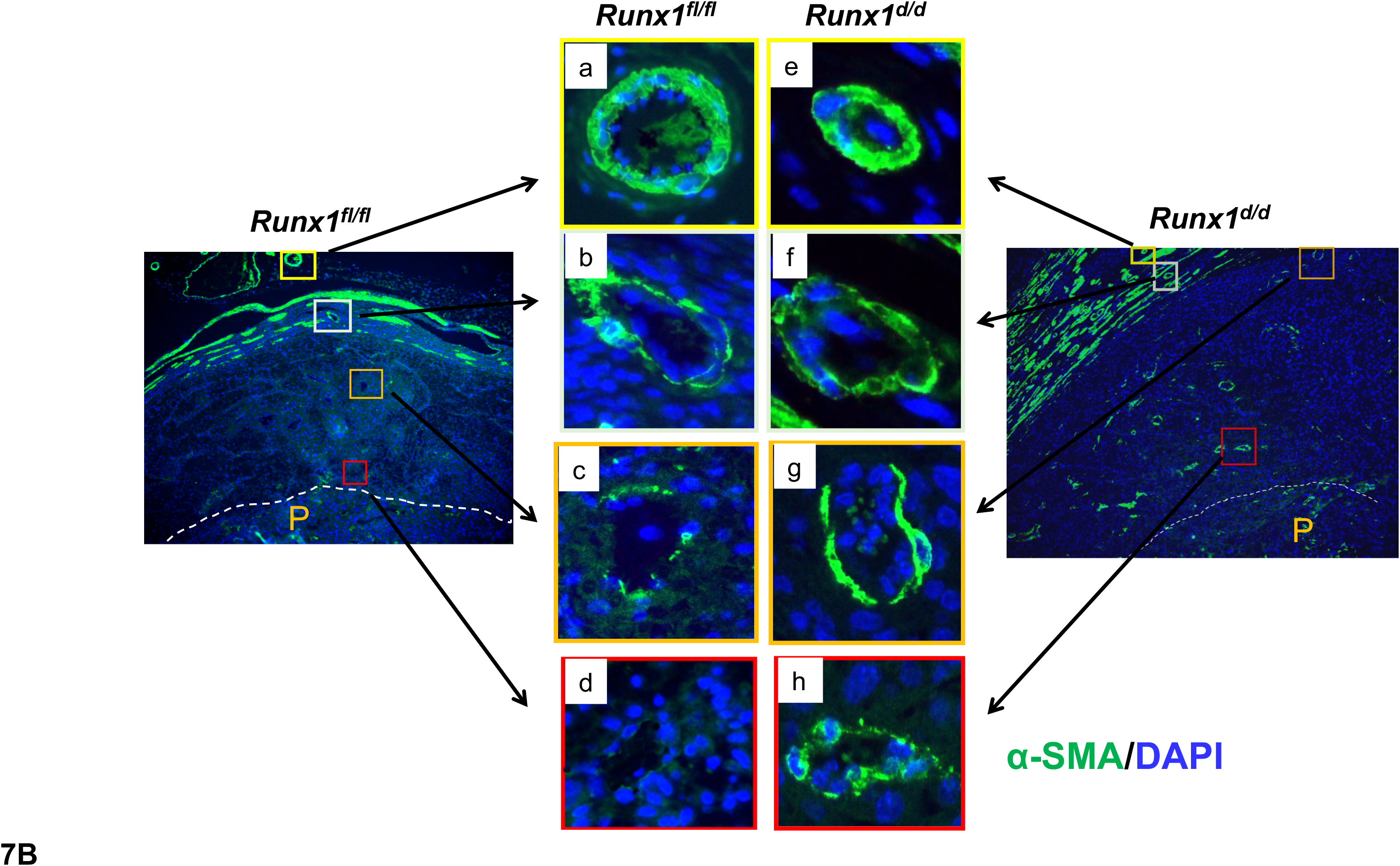
Impaired trophoblast invasion and spiral artery modification in *Runx1^d/d^* decidua. **A.** Uterine sections from *Runx1^f/f^* and *Runx1^d/d^* mice on day 12 of pregnancy were subjected to IF using KRT8 and *α*SMA antibodies. Trophoblast cells are indicated by cytokeratin 8 immunostaining (red) and spiral arteries are indicated by smooth muscle actin immunostaining (green). Boxed regions indicate magnified images (panels b and d). Spiral arteries (SA) are indicated by arrows. **B.** Colocalization of KRT8 and αSMA in day 12 uterine sections of *Runx1^fl/fl^* (a-d) and *Runx1^d/d^* (e-h) females. Boxed regions indicate magnified images. Panels a & e are radial arteries of *Runx1^fl/fl^* and *Runx1^d/d^* females. Panels b & f are spiral arteries in the myometrium, c & g are in distal endometrium, and d & h are in proximal endometrium. P denotes placenta. Dotted line separates maternal decidua from placental labyrinth. Data are representative images from n=3.

In Fig. 7B, we analyzed spiral arteries at various regions including myometrium, endometrial proximal and distal regions of the implantation sites by tracing α-SMA for arterial smooth muscle cells and cytokeratin 8 for trophoblasts. As expected, normal spiral artery modification was observed in *Runx1^fl/fl^* decidua, indicated by the disappearance of α-SMA from these vessels. By contrast, spiral arteries in *Runx1^d/d^* decidua retained their α-SMA expression, indicating a lack of dilation (Fig.7B). Taken together, our results indicated that decidual expression of Runx1 critically controls the differentiation of the trophoblast cells at the maternal-fetal interface and ensures development of proper vascular structure to support placental function.

### Runx1 controls the IGF signaling pathway

How does decidual Runx1 regulate trophoblast differentiation and placental development? To gain insights into the underlying mechanism, we analyzed the microarray-derived differentially expressed genes that are involved in cell proliferation and differentiation and found that the expression transcripts encoding two critical components of the insulin-like growth factor (IGF) signaling pathway, IGF2 and IGF binding protein 4 (IGFBP4), are markedly altered in the *Runx1^d/d^* decidua. While the levels of *Igf2* transcripts were sharply downregulated in the *Runx1^d/d^* decidua, we noted a marked upregulation of *Igfbp4* transcripts in *Runx1^d/d^* decidua on day 10 of pregnancy (Fig. 8A). Furthermore, ChIP analysis revealed that Runx1 binds directly to putative binding sites at the regulatory regions of *Igf2* and *Igfbp4* genes, indicating that it potentially regulates these genes directly (Fig. 8B). Immunohistochemical analysis of uterine sections on day 9 and day 10 of gestation, overlapping the time of trophoblast differentiation and placenta development, confirmed the downregulation of IGF2 protein and upregulation of IGFBP4 protein in *Runx1^d/d^* decidua (Fig. 8, C and D). Quantitation by Image J analysis estimated a ∼ 2 fold decrease in IGF2 expression and ∼ 2.5 fold increase in IGFBP4 protein expression in *Runx1*^d/d^ decidua.

**Figure 8.**
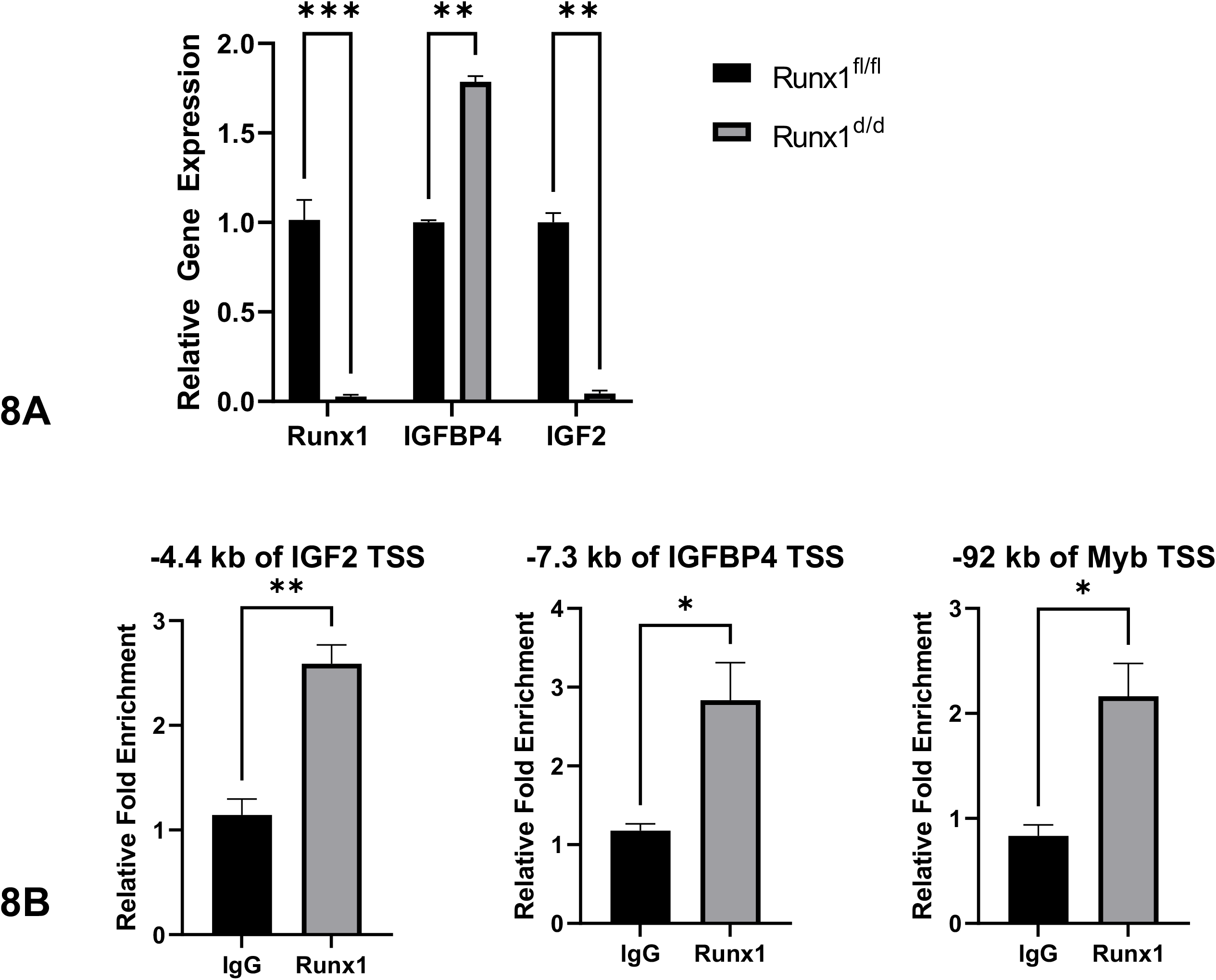
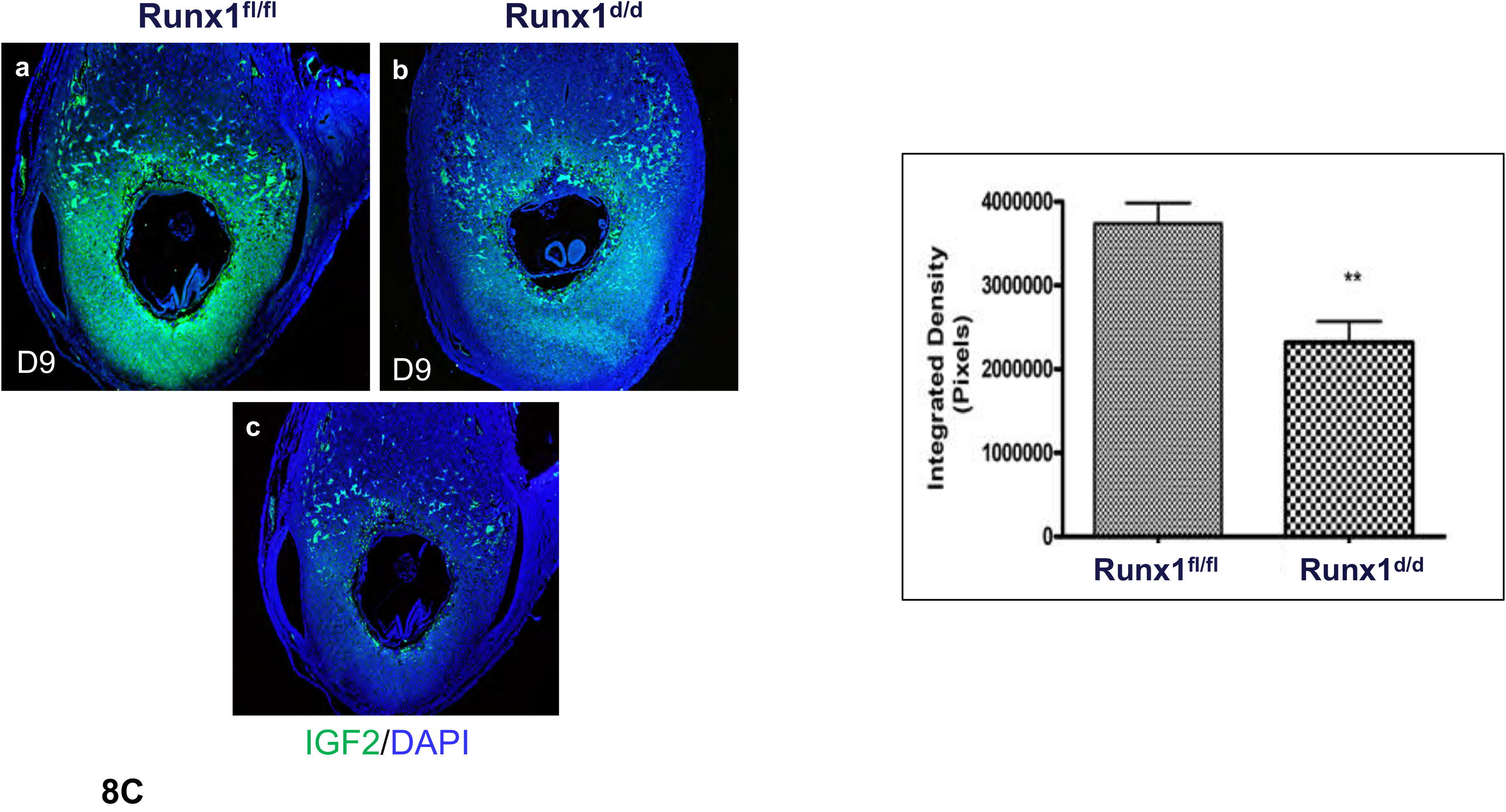
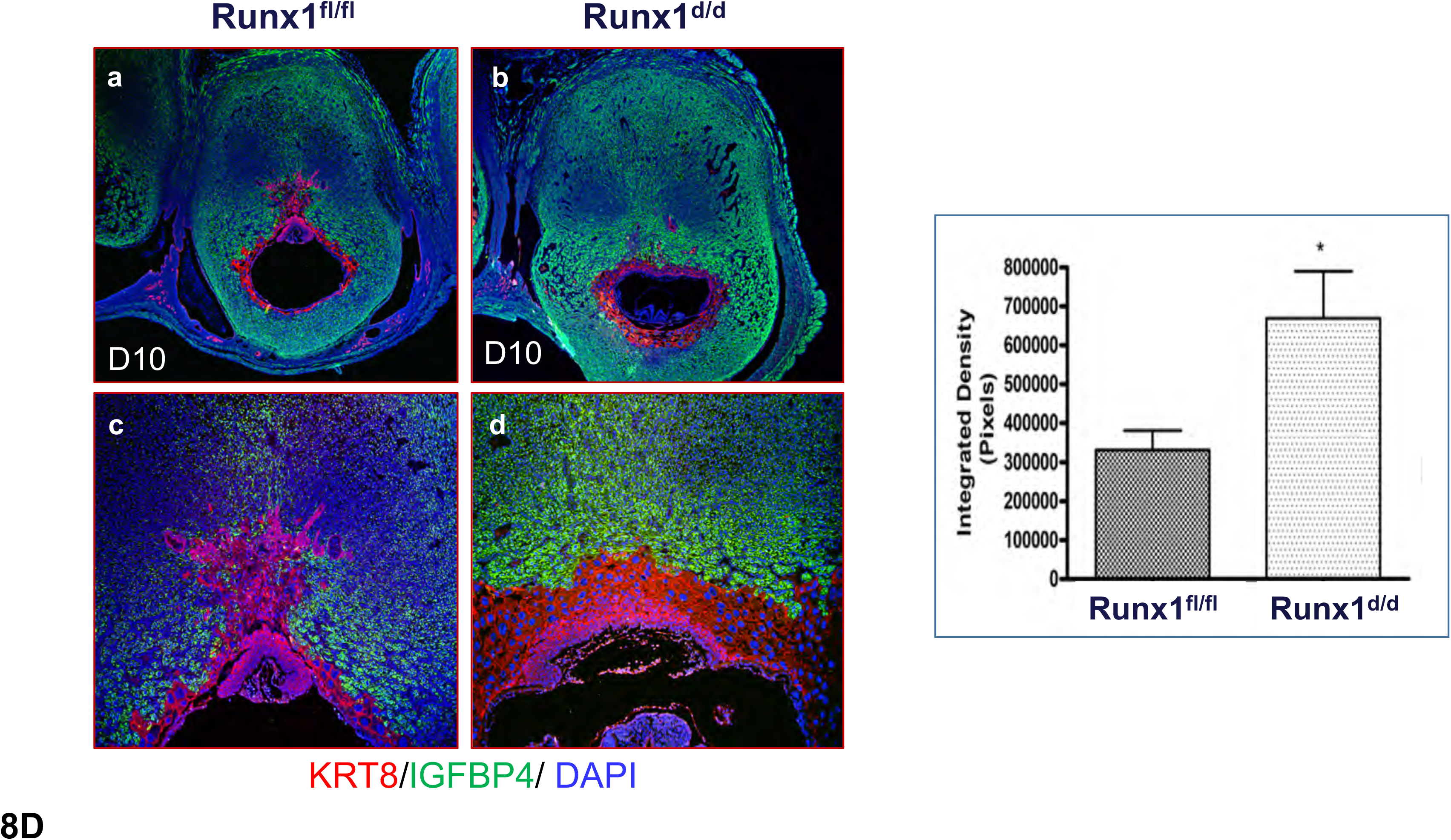
Runx1 regulates the gene expression of factors in the IGF-signaling pathway. **A.** RNA was extracted from mouse endometrial stromal cells from Runx1^fl/fl^ or Runx1^d/d^ mice that were isolated on day 4 of pregnancy and grown in culture for 48 h. Gene expression analysis was performed using primers specific for *Runx1*, *IGFBP4*, and *IGF2*. 36B4 was used to normalize gene expression. Data shown as mean fold change ± SEM (n=3), ** p<0.01, *** p<0.001, relative to Runx1^fl/fl^. **B.** ChIP was performed on mouse endometrial stromal cells isolated on day 4 of pregnancy and grown in culture with for 48 h. Chromatin enrichment was quantified by qPCR and normalized to a negative control locus. Primers flanking putative Runx1-binding sites upstream of IGF2 and IGFBP4 transcription start sites were used to quantify chromatin enrichment. Region – 92 kb of Myb TSS was included as a positive control locus. Data shown as mean fold change ± SEM (n = 3), * p<0.05, ** p<0.01, relative to IgG control. **C. Left:** Uterine sections of *Runx*1*^f/f^* (panel a) and *Runx*1*^d/d^* (panel b) mice on day 9 (n=3) were subjected to IF staining with IGF2 antibody. **Right:** Immuno-positive cells for IGF2 were analyzed by ImageJ software. Asterisks indicate statistically significant differences *p<0.05. Panel c indicates a negative control. **D. Left:** Dual IF staining of cytokeratin 8 (red) and IGFBP4 (green) are shown in *Runx1^f/f^* and *Runx1^d/d^* decidua on day 10 of pregnancy. Extensive trophoblast (red) invasion is evident in *Runx1*^f/f^ decidua. An elevated decidual IGFBP4 expression (green) is observed in *Runx1^d/d^* decidua (right). **Right:** Immuno-positive cells for IGFBP4 were analyzed by ImageJ software. Asterisks indicate statistically significant differences *p<0.05.

At the maternal-fetal interface, a critical balance of insulin-like growth factors, IGFs, and insulin-like growth factor binding proteins (IGFBPs), which curb the actions of these growth factors by reducing their local bioavailability, is thought to be an important determinant in the control of trophoblast differentiation that is essential for the development of proper functional placenta (28–30). The downregulation of IGF2 and concomitant abnormal accumulation of IGFBP4 in *Runx1*^d/d^ uterus results in a severely compromised IGF signaling at the maternal-fetal interface, hindering the proliferation and differentiation of trophoblast progenitor cells into invasive trophoblast cell lineages. Our results support the concept that the impaired trophoblast differentiation is a major contributor to the defective placentation observed in these mutant mice.

## DISCUSSION

Runx1 binds to the core consensus DNA sequence TGTGG to transcriptionally regulate downstream target gene expression (31). In the developing embryo, it critically controls pathways involved in the differentiation of hematopoietic progenitors into mature blood cells (31). The role of Runx1 in adult tissues remained largely unknown. Here, we report that Runx1 is induced in differentiating uterine stromal cells during the decidualization phase of pregnancy in mice. A previous study using *in vitro* cultures of mouse endometrial stromal cells suggested that Runx1 plays a role in decidualization (32). To address the *in vivo* function of this factor during embryo implantation and decidualization, we employed the *Runx1^d/d^* conditional null mice. Surprisingly, phenotypic analysis of *Runx1^d/d^* mice did not reveal any impairment in implantation or early phases of decidualization. Our study, however, revealed a novel role of this transcription factor in controlling multiple key events that guide the proper development of mouse placenta within a critical window during gestation days 9-15. Mice lacking *Runx1* in the decidua, therefore, present a unique model in which one can study the mechanisms by which maternal decidual factors regulate placenta development.

A major phenotypic consequence of the loss of Runx1 signaling in the decidua is a drastic decrease in the development of the uterine vascular network that supports embryonic growth. We found that Runx1 controls angiogenesis by regulating the expression of many downstream genes with known roles in this complex process. We, however, took special note of the reduced expression of Cx43, a gap junction protein (23). We had shown previously that conditional deletion of the *Cx43* gene using progesterone receptor (PGR)-Cre in *Cx43*^d/d^ mice did not affect the hypothalamic-pituitary-ovarian axis, implantation, or the myometrial function during early or mid-pregnancy, but led to an arrest of embryonic growth and widespread embryo resorption by gestation day 10. The stromal loss of Cx43, which disrupts gap junction communications between adjacent stromal cells during decidualization, resulted in drastically reduced expression of critical angiogenic factors, VEGFA, and angiopoietin, and a striking impairment in decidual angiogenesis (23). The uterine phenotype of the *Cx43*^d/d^ mice is, therefore, remarkably similar to that of *Runx1*^d/d^ mice. This finding is supported by our demonstration that Runx1 directly regulates the expression of the *Cx43* gene. Loss of Cx43 expression in *Runx1*^d/d^ uteri, disrupts gap junctional signaling that presumably controls the expression of downstream angiogenic factors, such as VEGFA and angiopoietin 2, by decidual cells via yet unknown mechanisms.

Another major finding of this study is that Runx1-directed mechanisms in the decidual cells that control trophoblast cell lineage differentiation and function. In mice, around days 7.5-8 of gestation, trophoblast progenitor cells of the ectoplacental cone (EPC) of the developing placenta proliferate and differentiate into distinct trophoblast cell lineages, such as secondary TGCs and spongiotrophoblasts, which play essential roles at the maternal-fetal interface (24). By days 10-12 of gestation, maternal decidua is invaded by endovascular TGCs and spongiotrophoblasts. We noted that trophoblast differentiation and invasion into the decidua are curtailed in the absence of Runx1. Specifically, the differentiated spongiotrophoblasts could not reach the spiral arteries in *Runx1*^d/d^ decidua and did not displace the smooth muscle actin layer in these arteries. Additionally, population of spongiotrophoblasts and their invasion into *Runx1*^d/d^ decidua were markedly reduced on day 10 of gestation. Since the *Runx1* conditional allele is intact in the embryos present in pregnant *Runx1*^d/d^ females, any defect in trophoblast differentiation and migration into the decidua is clearly a consequence of aberrant maternal signals originating in the *Runx1*^d/d^ decidua.

It is generally thought that autocrine secretion of IGFs by the embryo drives trophoblast proliferation and differentiation (30, 33, 34). It is well-documented that the trophoblasts express IGF2 (28, 30, 35). Interestingly, we observed that decidual stromal cells also express significant expression of IGF2 mRNA and protein. This raises the possibility that the decidual IGF2 may act in a paracrine manner to promote trophoblast proliferation and differentiation. On the other hand, maternal decidua modulates the actions of IGFs by secreting IGFBPs, which control IGF bioavailability (36, 37). Two members of the IGFBP family, IGFBP1 and IGFBP4, were previously reported to be expressed in the mouse uterus during early pregnancy (38). While IGFBP1 was expressed in mouse uterine epithelial cells, IGFBP-4 is the predominant IGFBP in stromal cells during the decidual phase of pregnancy (38). Clinically, increased IGFBP4 level in the maternal circulation in early pregnancy was previously linked to development of intrauterine growth restriction (37, 39). Interestingly, our data showed abnormally elevated levels of IGFBP4 in *Runx1^d/d^* decidua compared to the *Runx1^f/f^* sample. These results support the concept that combined effects of reduced expression of decidual IGF2 and abnormal accumulation of IGFBP4 at the maternal-fetal interface in *Runx1^d/d^* uterus result in suppression of IGF signaling, hindering the proliferation and differentiation of trophoblast progenitor cells into invasive trophoblast cell lineages.

In conclusion, our study reveals a set of molecular, cellular, and integrative mechanisms that are dictated by Runx1 in the decidua and ensure coordination of the endometrial differentiation and angiogenesis with embryonic growth within a critical time window when the placenta begins to form in pregnant mice. Apart from the *Runx1*^d/d^ model described in this study, two other animal models including conditional deletion of Rac1 and bone morphogenetic protein receptor type 2 (BMPR2) in the uterine decidua have addressed the mechanisms by which maternal factors control placenta development (40, 41). While there are phenotypic similarities such as abnormal vascular development and impaired placental function among *Runx1*^d/d^, *Rac1^d/d^* and *Bmpr^d/d^* mouse models, it remains to be determined whether Runx1, Rac1, and BMPR2 pathways converge, or they function via distinct mechanisms to control placental development. It also remains to be addressed whether Runx1 pathway is conserved in human endometrium. Further studies using human endometrial specimens from normal cycling women and patients with recurrent pregnancy loss may reveal whether an impaired Runx1 pathway is linked to dysregulated endometrial gene expression, contributing to disrupted maternal-embryo coordination, affecting establishment of pregnancy, and causing miscarriage.

## ACKNOWLEDGMENTS

This work was supported by the Eunice Kennedy Shriver NICHD/NIH U54 HD 055787 and R01 HD090066 (to ICB and MKB). JRB is supported by the NIEHS training grant T32 ES007326. We thank Dr. Shanmugasundaram Nallasamy for generating and maintaining the Runx1^f/f^ and Runx1^d/d^ mice.

## MATERIALS AND METHODS

### Animals

Mice were maintained in the animal facility at the University of Illinois, College of Veterinary Medicine, in accordance to the institutional guidelines for the care and use of laboratory animals and in accordance with the National Institutes of Health standards for the use and care of animals. Mice were housed in an animal room with temperature of 22 °C and 12L:12D cycles. Food and water were provided ad libitum. The Institutional Animal Use and Care Committee at the University of Illinois at Urbana-Champaign approved all procedures involving animal care, euthanasia, and tissue collection.

Conditional *Runx1*-null mice (*Runx1^d/d^*) were generated by crossing mice harboring a ‘floxed’ *Runx1* gene termed as *Runx1^f/f^* with *Pgr-Cre* knock-in mice. The *Runx1* ‘floxed’ mice were kindly provided by Dr. Gary Gilliland (42). Drs. Francesco J. DeMayo and John P. Lydon provided the *Pgr-Cre* knock-in mice (19). This strategy has been used extensively to ablate ‘floxed’ genes in tissues expressing PGR (43, 44).

### Fertility assessments, timed pregnancies, and tissue collection

To test fertility, *Runx1^f/f^* and *Runx1^d/d^* mice of reproductive age (7-8 weeks) were paired with fertile wild-type males for six months. The total number of pups born in each litter and the number of pregnancies during this period was recorded.

For experiments involving timed pregnancies, female mice were mated with adult wild-type males of known fertility. For tissue collection, all animals were euthanized by CO_2_ asphyxiation. Uteri and ovaries were collected at different time points during pregnancy and the tissues were immersion-fixed in 10% (vol/vol) neutral-buffered formalin (NBF) for histological evaluation or flash frozen in liquid N_2_ for RNA isolation or frozen sectioning. As a reference for our experiments, the identification of a copulatory plug indicated day 1 of pregnancy.

### Serum hormone assay

Following euthanasia, blood was drawn via cardiac puncture using a 30-gauge needle and transferred into a sterile 1.5 mL tube. The blood samples were incubated at room temperature for 90 min to allow clot formation. After the incubation, the clot was removed with a sterile pipette tip and the samples were spun at 2000 x *g* for 15 min at room temperature. The serum samples were transferred into a new sterile 1.5 mL tube and stored at -80 °C until analyzed. Serum hormones were measured by radioimmunoassay at the Ligand Core facility, University of Virginia, Charlottesville.

### Alkaline phosphatase activity

Alkaline phosphatase (ALPL) activity was detected following previously published protocols (21), with modifications. Briefly, frozen uterine sections were fixed in 10% NBF for 10 min, and then washed with 1x phosphate-buffered saline (PBS) three times for 5 min each. The uterine sections were then incubated in the dark at 37 °C for 30 min in a solution containing 0.5 mM naphthol AS-MX phosphate (ALPL substrate) and 1.5 mM Fast Blue RR in 0.1 M Tris-HCl, pH 8.5. Alkaline phosphatase activity releases orthophosphate and naphthol derivatives from the ALPL substrate. The naphthol derivatives are simultaneously coupled with the diazonium salt (Fast Blue RR) to form a dark dye marking the site of enzyme action. The slides were rinsed in tap water to terminate the enzymatic reaction. Stained uterine sections were visualized under an Olympus BX51 microscope equipped for light imaging and connected to a Jenoptik ProgRes C14 digital camera with c-mount interface containing a 1.4 Megapixel CCD sensor.

### Immunohistochemistry (IHC)

Uterine tissues were processed and subjected to immunohistochemistry (IHC) as described previously (40, 45). Briefly, paraffin-embedded tissues were sectioned at 5 µm and mounted on microscopic slides. Sections were deparaffinized in xylene, rehydrated through a series of ethanol washes, and rinsed in water. Antigen retrieval was performed by immersing the slides in 0.1M citrate buffer solution, pH 6.0, followed by microwave heating for 25 min. The slides were allowed to cool and endogenous peroxidase activity was blocked by incubating sections in 0.3% hydrogen peroxide in methanol for 15 min at room temperature. After washing with PBS for 15 min, the slides were incubated in a blocking solution for 1 h. This was followed by incubation overnight at 4°C with antibodies specific for RUNX1 (Santa Cruz, SC-28679, 1;100), insulin-like growth factor-binding protein 4 (IGFPB4, 1:200, Novus, NBP1-80549), insulin growth factor 2 (Abcam, ab262713, 1:500).

Placental sections were also subjected to immunohistochemistry using the primary antibodies: cytokeratin 8 (KRT8, 1:50, Developmental Studies Hybridoma Bank, TROMA-1), platelet/endothelial cell adhesion molecule 1 (PECAM1/CD31, 1:250, Abcam, ab124432), placental lactogen 1 (PL1, 1:200, Santa Cruz Biotechnology, SC-34713), PLF (Santa Cruz, SC-47345, 1:100), smooth muscle actin (SMA, Abcam, ab5694, 1:300). Secondary antibodies including fluorescent-tagged rhodamine donkey anti-rabbit, 488 donkey anti-mouse, 488 donkey anti-rabbit, 488 donkey anti-goat were purchased from Jackson Immuno Research. Fluoromount-G with DAPI was purchased from eBiosciences. Pictures were taken using the Olympus BX51 microscope equipped for fluorescent imaging and connected to a Jenoptik ProgRes C14 digital camera with c-mount interface containing a 1.4 Megapixel CCD sensor. Fluorescent images were processed and merged using Adobe Photoshop Extended CS6 (Adobe Systems).

### Quantitative Real time PCR analysis (qPCR)

Total RNA was isolated from uteri, ovaries, and cells using a standard TRIzol-based protocol. The RNA concentration of each sample was determined at 260 nm using a Nanodrop ND1000 UV-Vis spectrophotometer (Nanodrop Technologies). RNA samples were reverse transcribed using the High Capacity cDNA Reverse Transcription kit (Applied Biosystems) according to the manufacturer’s instructions. Primers specific for genes of interest were developed and real-time quantitative PCR (qPCR) reactions were carried out using SYBR-green master mix (Applied Biosystems) in a 7500 Applied Biosystems Real-time PCR machine (Applied Biosystems). For each sample, the mean threshold cycle (Ct) was calculated from Ct values obtained from three replicates. The normalized ΔCt in each sample was calculated as mean Ct of target gene subtracted by the mean Ct of the reference gene. ΔΔCt was then calculated as the difference between the ΔCt values of the control and mutant samples. The fold change of gene expression in each sample relative to a control was generated using the 2^−ΔΔCt^ mathematical model for relative quantification of quantitative PCR (40, 45). The mean fold induction and SEM were calculated from at least three or more independent experiments. The housekeeping gene *RPLP0* (*36B4*), which encodes a ribosomal protein, was used as a reference gene. Reported data consists of mean fold induction ± SEM.

### Chromatin immunoprecipitation (ChIP) analysis

ChIP assays were performed using the EZ-ChIP kit (Millipore) according to the manufacturer’s instructions. Briefly, mouse endometrial stromal cells were fixed with 1% formaldehyde for 10 min and excess formaldehyde was quenched with 0.25 M glycine for 5 min. Cells were then washed twice with cold PBS, scraped, and collected in PBS containing protease inhibitors in a conical tube and centrifuged. Cell pellets were resuspended in lysis buffer containing protease inhibitors for 15 min to lyse the cells. Next, chromatin was sonicated in three 15-s pulses with cooling between pulses (Fisher Scientific model 100 Sonic Dismembrator). One percent of the cell lysate was used for input control, and the rest was used for immunoprecipitation using an antibody against Runx1. After incubation at 4 C overnight, protein G-agarose beads were used to isolate the immune complexes. Complexes were washed with low salt, high salt, LiCl buffer, twice with TE buffer, and then eluted from the beads and heated at 65°C for 6 hours to reverse the cross-linking. After digestion with RNAase A and proteinase K, DNA fragments were purified using QIAquick PCR purification kit (Qiagen). For ChIP, real-time quantitative PCR primers were designed to amplify the potential Runx1 binding sites in the Connexin 43, IGF2, IGFBP4, or Myb promoter regions. The resulting signals were normalized to input DNA.

### Microarray and expression analysis

Mouse endometrial stromal cells (MESC) were isolated from Runx1f/f and Runx d/d female mice on day 10 of pregnancy. Total RNA was extracted from MESC using a standard TRIzol-based protocol. RNA integrity was verified using Agilent 2100 bioanalyser (Agilent Technologies Inc., Santa Clara, CA, USA) at the Biotechnology Center of the University of Illinois, Urbana and Champaign. Each RNA sample was processed for microarray hybridization using Affymetrix GeneChip Mouse Genome 430A 2.0 array, which contains probes that represented approximately 14,000 annotated gene sequences, following the established protocol. Microarray data reported in this paper is deposited in the Harvard Dataverse. https://doi.org/10.7910/DVN/GOTVYC

### Statistical analyses

Statistical analyses were performed as we have done previously (40). Experimental data were collected from a minimum of four independent samples, which were subjected to the same experimental conditions. All numerical data are expressed as mean ± SEM. Statistical analysis was done using a two-tailed Student’s *t-test*. One-way ANOVA with a Tukey post hoc test was performed in studies involving multiple comparisons. An analysis of equal variances was done on all numerical data to determine whether a parametric or non-parametric hypothesis test was appropriate. Data were considered statistically significant at p ≤ 0.05 and are indicated by asterisks in the figures. All data were analyzed and plotted using GraphPad Prism 9.0 (GraphPad Software).

---

**Figure. S1:**
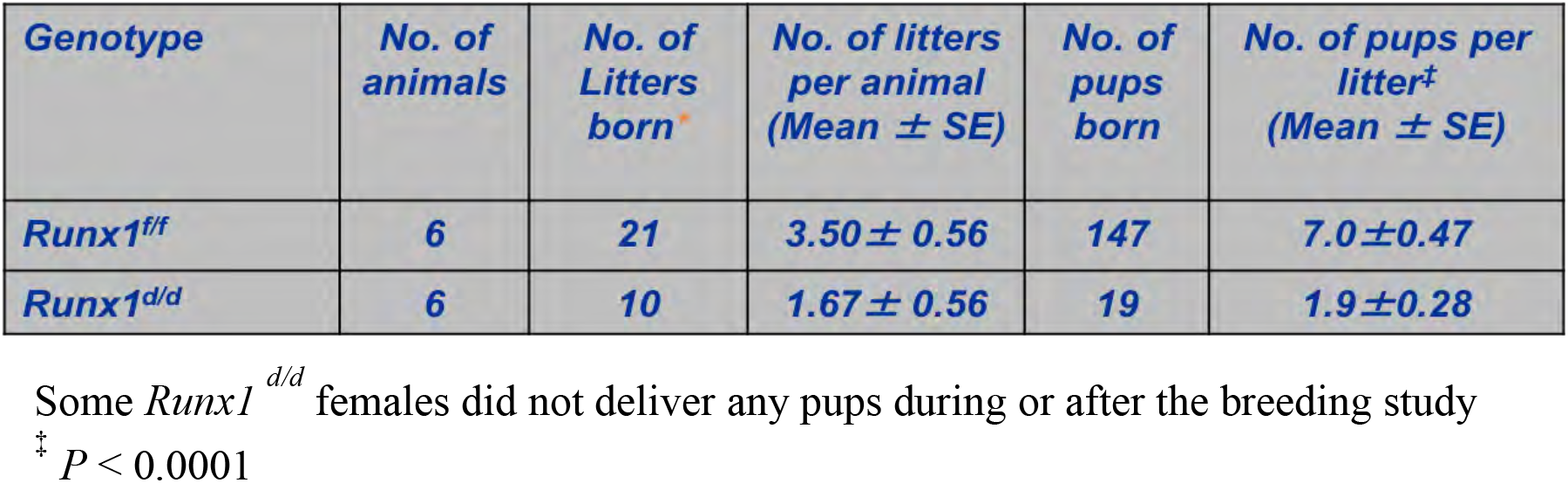
Ablation of uterine *Runx1* leads to severe female infertility. The results of a breeding study are shown.

**Figure S2.**
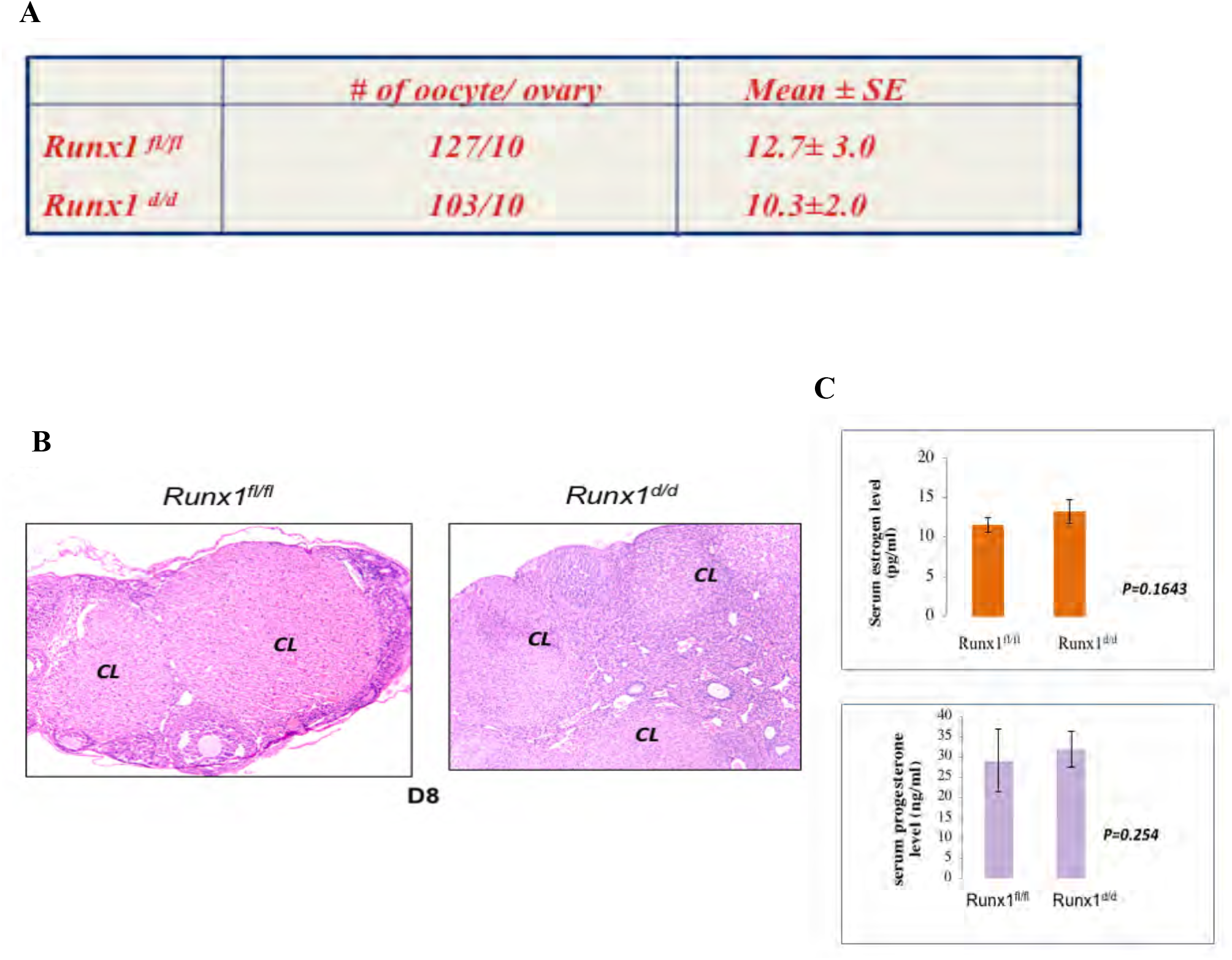
Ovarian function remains unaffected in *Runx1^d/d^* mice. **A.** Age-matched prepubertal *Runx1^f/f^* (n=6) and *Runx1^d/d^* mice (n=6) were subjected to superovulation. The oocytes were recovered and counted at 18h after hCG administration (values are mean ± SEM). **B.** H & E staining of ovarian sections from *Runx1^f/f^* and *Runx1^d/d^* mice on day 8 of pregnancy. CL indicate corpora lutea. **C.** Progesterone and Estrogen levels in serum of *Runx1^f/f^* (n=6) and *Runx1^d/d^* (n=6) mice on day 8 of pregnancy. Values are represented as means ± SEM.

## Notes

### Competing Interest Statement

The authors have declared no competing interest.

